# Rational design and adaptive management of combination therapies for Hepatitis C virus infection

**DOI:** 10.1101/008466

**Authors:** Ruian Ke, Claude Loverdo, Hangfei Qi, Ren Sun, James O. Lloyd-Smith

## Abstract

Recent discoveries of direct acting antivirals against Hepatitis C virus (HCV) have raised hopes of effective treatment via combination therapies. Yet rapid evolution and high diversity of HCV populations, combined with the reality of suboptimal treatment adherence, make drug resistance a clinical and public health concern. We develop a general model incorporating viral dynamics and pharmacokinetics/pharmacodynamics to assess how suboptimal adherence affects resistance development and clinical outcomes. We derive design principles and adaptive treatment strategies, identifying a high-risk period when missing doses is particularly risky for *de novo* resistance, and quantifying the number of additional doses needed to compensate when doses are missed. Using data from large-scale resistance assays, we demonstrate that the risk of resistance can be reduced substantially by applying these principles to a combination therapy of daclatasvir and asunaprevir. By providing a mechanistic framework to link patient characteristics to the risk of resistance, these findings show the potential of rational treatment design.

## Introduction

Hepatitis C virus (HCV) affects approximately 170 million people and chronic infections can lead to cirrhosis and hepatocellular carcinoma (Lavanchy 2009; Thomas 2013). Recently, development of direct acting antivirals (DAAs) against HCV infection has revolutionized the field of HCV treatment, because of their high potency, broad applicability and mild side effects (Gane 2011; Scheel and Rice 2013). Combination therapies of DAAs have achieved remarkably high rates of sustained virological response in clinical trials (Lok, Gardiner et al. 2012; Pol, Ghalib et al. 2012; Afdhal, Zeuzem et al. 2014; Feld, Kowdley et al. 2014; Kowdley, Gordon et al. 2014; Zeuzem, Jacobson et al. 2014). However, due to the relatively low genetic barriers of most DAAs (Pawlotsky 2011; Robinson, Tian et al. 2011; Qi, Olson et al. 2014), the high intrinsic mutation rate of HCV (Powdrill, Tchesnokov et al. 2011; Ribeiro, Li et al. 2012), and the high viral diversity (Pybus, Charleston et al. 2001; Simmonds 2004; Thomas 2013), combined with the reality of suboptimal treatment adherence (Lo Re, Teal et al. 2011; Lieveld, van Vlerken et al. 2013), viral resistance represents a clinical and public health concern (Sarrazin and Zeuzem 2010; Pawlotsky 2011). Indeed, resistance to single DAAs has already been observed frequently for many candidate DAAs, and patients must be treated with combination therapies to avoid treatment failure. If not properly managed, resistance could quickly develop to combination therapies and render these new DAAs useless, as observed for other antimicrobial treatments, squandering the potential health gains from these recent breakthroughs (DiMasi, Hansen et al. 2003; Roberts, Hota et al. 2009; Smith, Okano et al. 2010).

Suboptimal patient adherence to dosing regimens is a crucial risk factor for resistance development in both HIV and HCV treatments (Paterson, Swindells et al. 2000; Bangsberg, Perry et al. 2001; Lo Re, Teal et al. 2011; Lieveld, van Vlerken et al. 2013). Although high rates of sustained virological response have been achieved in clinical trials, adherence levels may vary substantially among the vast population of infected patients, owing to long treatment periods and complicated regimens associated with DAA combination therapies (Weiss, Brau et al. 2009; Lo Re, Teal et al. 2011; Evon, Esserman et al. 2013; Gordon, Yoshida et al. 2013; Lieveld, van Vlerken et al. 2013). Rational design of combination therapy regimens, enabling individualized regimens based on the genetic composition of a patient’s infection and real-time adjustments for missed doses, is a top research priority to avoid resistance (Spellberg, Guidos et al. 2008; Gelman and Glenn 2010; zur Wiesch, Kouyos et al. 2011; Lieveld, van Vlerken et al. 2013). Mathematical models are well suited to address this problem. Previous modeling studies for HIV infections have illuminated potential mechanisms underlying treatment failure and explained puzzling clinical observations (Wahl and Nowak 2000; Rosenbloom, Hill et al. 2012). However, HCV is a curable disease and its infection, goal of treatment and mechanism of resistance differ from HIV in many respects (Soriano, Perelson et al. 2008), including no known latent reservoir and a finite treatment period to eradicate the virus. Here, by integrating pharmacokinetics/pharmacodynamics (PK/PD) and viral dynamics into mathematical models, we develop the first general theory to assess the impacts of suboptimal adherence on the outcome of DAA-based therapies for HCV infection. We derive design principles that can be generalized to therapies involving different classes and different numbers of drugs. Using large-scale data from in vitro resistance assays and human clinical trials, we apply this framework to a combination therapy of daclatasvir and asunaprevir (Suzuki, Ikeda et al. 2012), and derive evidence-based adaptive treatment strategies for treatment protocols over time according to resistance profiles and adherence patterns.

## Results

Resistance to antiviral treatments can develop through selection of preexisting mutants or *de novo* generation of new mutants. A core principle for designing effective combination therapy is that, if patients fully adhere to the treatment regimen, the treatment must suppress all preexisting mutants and *de novo* resistance should be unlikely (Ribeiro and Bonhoeffer 2000). Missing doses, however, can lead to suboptimal drug concentrations, allowing growth of some preexisting mutants with partially resistant phenotypes. Growth of these mutants allows the viral population to survive longer, possibly generating further mutations that contribute *de novo* resistance against the full combination therapy. For example, consider a combination therapy of two DAAs, A and B, as shown in Fig. 1A. If missed doses and pharmacokinetics lead to a drop in the concentration of drug A, this allows growth of the preexisting mutant, AB’, (which is already resistant to drug B), thus opening opportunities to generate the fully resistant mutant, A’B’. Therefore, the dynamics of the subset of preexisting mutants that have a high level of resistance against single DAAs determine resistance evolution and treatment outcomes for combination therapies. In the following, we denote these mutants as ‘*partially resistant’ mutants*.

**Figure 1.**
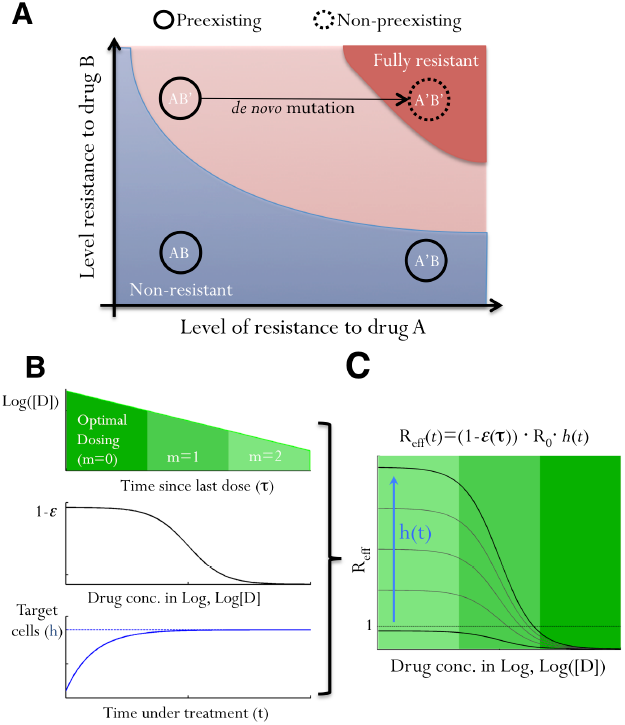
The impacts of suboptimal adherence on viral fitness. **(A)** A schematic illustrating how a non-preexisting mutant, A’B’, fully resistant to a combination therapy involving two drugs, A and B, can be generated when adherence is suboptimal. Each black circle represents a mutant on the parameter space of resistance levels to A and B. AB, A’B and AB’ are preexisting mutants that are non-resistant, resistant to A only and resistant to B only, respectively. Colored areas denote parameter regimes where mutants are fully resistant to the therapy (red), can grow if doses are missed (pink), and do not grow (blue). Note that the pink area can grow or shrink on the parameter space depending on the number of consecutively missed doses and drug PK/PD, and mutants lying in the pink area are ‘partially resistant mutants’. **(B)** The dynamics of viral strains under treatment are determined by several factors: drug concentration, [D], which decreases with an increasing number of missed doses, *m* (upper panel); how viral replication is affected by drug (1-*ε*; middle panel); and the relative number of target cells, *h*(*t*) (lower panel). Upon effective treatment, *h*(*t*) increases to the infection-free level. **(C)** We integrate all these factors into a single fitness parameter, *R_eff_*(*t*). Viral fitness increases as drug concentration drops (indicated by shades of green) and as target cell abundance rises (the blue arrow). Values of *R_eff_*(*t*) can exceed 1, i.e. positive growth, if doses are missed after a period of effective treatment.

### The effective viral fitness, R_eff_(t)

The fitness of a particular strain in a treated patient is determined by the PK/PD of the drug, the level of resistance of the strain, and the availability of target cells, i.e. uninfected hepatocytes for HCV (Fig. 1B). We can integrate all these factors (for any class of DAA therapy) into a single number, the effective reproductive number under treatment, *R_eff_*(*t*) (Fig. 1C). *R_eff_*(*t*) is the average number of cells infected by viruses produced by a single infected cell. It acts as a measure of viral fitness, and can be calculated as:

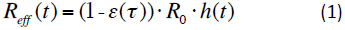

where *t* is time since treatment starts, *τ* is the time since last dose, *ε*(*τ*) is the efficacy of the drug combination at time τ during the dosing cycle, *R*_0_ is the reproductive number of the virus in the absence of treatment, and *h*(*t*) is the normalized abundance of target cells (see Supplementary Materials). Under effective treatment, the availability of target cells, *h*(*t*), increases quickly to reach the infection-free level (Rong, Dahari et al. 2010), and therefore, the overall viral fitness increases over time as *h*(*t*) increases under effective treatment (Fig. 1B,C). When adherence is optimal, the value of *R_eff_* for a partially resistant mutant is always less than 1 (i.e. viral suppression); however, if doses are missed, drug concentration declines exponentially and *R_eff_*can become greater than 1 (i.e. viral growth) (Fig. 1C).

### The growth of partially resistant mutants and the need for extended treatment

We now consider how suboptimal adherence impacts the dynamics of partially resistant mutants. As an illustration, we contrast simulations assuming perfect adherence versus suboptimal adherence. Missing doses leads to rapid decreases in drug concentration, and thus increases in R*_eff_* of a partially resistant mutant (Fig. 2A-C). This means that extra doses are needed to compensate for the missed doses to suppress the mutant to extinction (Fig. 2D), and also that the number of newly infected cells rises substantially, which increases the opportunity for *de novo* resistance (Fig. 2E).

**Figure 2.**
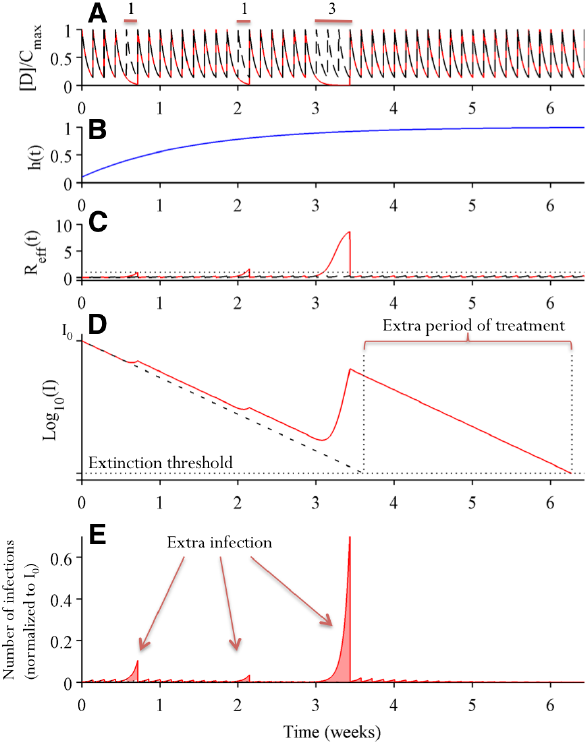
Suboptimal adherence prolongs treatment time needed to eliminate *partially resistant mutants* and increases the risk of *de novo* evolution of fully resistant strains. Two simulations assuming perfect adherence (dashed black lines) and imperfect adherence (solid red lines) are shown. In the simulation assuming imperfect adherence, single doses are missed at day 5 and 15, and 3 consecutive doses are missed during days 22-24. **(A)** Drug concentration over time normalized by the maximum drug concentration *C_max_*. **(B)** The abundance of target hepatocytes, which rebounds after the initiation of combination therapy. **(C)** Viral fitness of the partially resistant mutant under consideration. Missing doses increases the value of *R_eff_*, especially when multiple consecutive doses are missed or when *h*(*t*) has increased to high levels. **(D)** The dynamics of cells infected by the PM (on log_10_ scale). The number of infected cells declines almost exponentially when doses are taken. Missed doses allow the number of infected cells to rebound. This means that an additional period of treatment is needed to suppress the mutant below the extinction threshold level, i.e. to achieve viral elimination. **(E)** The number of cells newly infected by the partially resistant mutants. Missed doses lead to substantial numbers of additional new infections, especially when 3 consecutive doses are missed.

We approximate the time-varying values of *R_eff_*(*t*) during periods when doses are missed, by calculating the average effective reproductive number, *R_ave,m_*, as (see Materials and Methods):

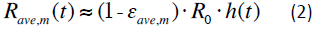

where *t* is the time when the patient starts to miss doses, *m* is the number of consecutive doses missed and *ε_ave,m_* is the average drug inhibition during the period when *m* consecutive doses are missed. This allows us to generalize our theory to any DAA combinations for which *ε_ave,m_* can be either estimated from pharmacokinetics/pharmacodynamics data or calculated from mutant resistance profiles (Wahl and Nowak 2000).

We then ask, if *m* consecutive doses are missed beginning at time *t*, how many extra doses, *N_m_*, are needed to compensate? This number, which we denote ‘compensatory doses’, can be approximated as (see Materials and Methods):

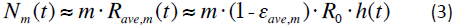

This allows us to estimate the total duration of treatment needed to clear infection for a given adherence pattern. Furthermore, since *h*(*t*) increases over time under effective treatment (Rong, Dahari et al. 2010), Eqn. 3 shows that a higher number of extra doses are needed to eliminate the infection if doses are missed later in treatment.

### De novo generation of fully resistant mutants

To assess the risk that a partially resistant lineage will give rise to full resistance, we calculate the expected number of target cells, Φ*_m_*, that become infected by fully resistant mutant viruses due to *de novo* mutation during a period when *m* consecutive doses are missed. This quantity is the product of the cumulative number of cells newly infected by a partially resistant mutant and the effective mutation rate from that mutant to the fully resistant mutant, *µ_eff_* (see Materials and Methods):

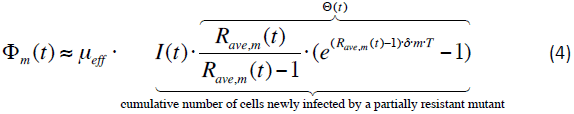

where *I*(*t*) is the number of cells infected by the partially resistant mutant at time *t* when the first dose is missed, and Θ(*t*) represents the potential to generate new infections. *δ* is the death rate of infected hepatocytes, and *T* is the scheduled interval between two doses. Φ*_m_* quantifies the risk that a fully resistant mutant infects target cells, but whether it emerges and becomes established within the host depends on its fitness and the stochastic dynamics of invasion (Alexander and Bonhoeffer 2012; Loverdo, Park et al. 2012; Loverdo and Lloyd-Smith 2013).

The strong dependence of Φ*_m_* on *µ_eff_* predicts that designing combination therapies to increase the genetic barrier to full resistance, e.g. using DAAs with higher genetic barrier or adding an extra drug into the combination, can reduce Φ*_m_*by orders of magnitude or more, thus it would lead to drastic reductions in the probability of generating full resistance (compare trajectories a and b in Fig. 3A).

**Figure 3.**
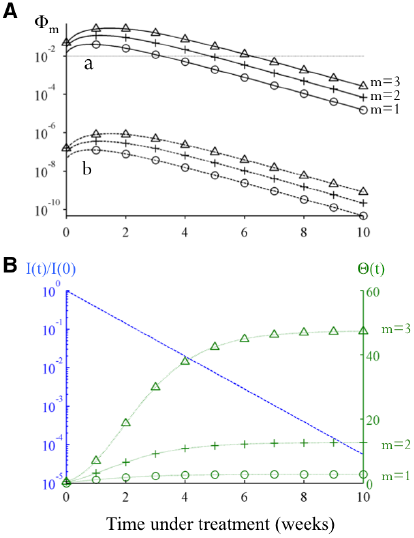
There is a high-risk window early in treatment when missing doses is more likely to cause *de novo* resistance. **(A)** The changes in the risk of *de novo* resistance, Φ_m_, generated by a partially resistant mutant over time. The two sets of trajectories, A and B, differ in that the value of *μeff* for trajectory B is smaller by a factor of 10-5 (representing one additional nucleotide mutation) than the value set for trajectory A. Each set of trajectories shows the risk when the number of doses missed (*m*) is 1,2 or 3. **(B)** Dynamics of the two time-varying quantities in Eqn.4, i.e. the number of cells infected by the partially resistant mutant relative to the initial number before treatment (I(t)/I(0); blue dashed line), and the value of Θ(*t*), green dotted lines, as shown in Eqn.4. Under effective treatment, the number of infected cells *I*(*t*) decreases exponentially, while the number of target cells rebounds to the infection-free level quickly, causing an increase in R*_ave,m_* and thus Θ(*t*). Together these changes cause Φ*_m_* to increase initially and then to decrease exponentially at longer times (as seen in panel A).

Eqn. 4 also allows us to assess when during treatment it is most risky to miss doses, which can inform treatment guidelines. Changes in two quantities, *I*(*t*) and Θ(*t*), determine changes in Φ*_m_* over the course of a treatment regimen. For as long as adherence is perfect, *I*(*t*) decreases exponentially, while Θ(*t*) increases over time since *R_ave,m_*(*t*) increases as the abundance of target cells rises over time (Fig. 3B). Thus the value of Φ*_m_* first increases (due to rapid recovery of target cells) and then decreases exponentially (due to decrease of infected cells). This leads to a high-risk window period, during which missing doses is especially risky for generating full resistance (Fig. 3A). This qualitative finding is robust to changes in model parameters, though quantitative predictions of the risk of full resistance depend on the fitness of the mutant (*R*_0_), the half-life of infected cells (*δ*), and the rate at which the target hepatocytes become available upon treatment (Fig.S1).

### Design principles and adaptive treatment strategy for DAA combination therapy

These results suggest principles for rational optimization of treatment outcomes. Individualized therapies could be designed for patients with risk factors for low adherence, by selecting drug combinations that impose a higher genetic barrier than required to suppress all preexisting mutants, to reduce the risk of *de novo* resistance.

*Adaptive treatment strategies* could be developed based on the theoretical findings shown above. For a particular combination therapy, the high-risk window period for missing doses can be calculated by integrating the values of Φ*_m_* for all partially resistant mutants present in a patient. Then, for patients with risk of low adherence, supervised dosing during the high-risk window period would reduce the risk of resistance and treatment failure. Another alternative is to treat the patient using a higher number of DAAs in combination during the high-risk period, and then switch back to a combination with a lower number of DAAs afterwards. If doses are missed during treatment, the patient should be treated with extra doses, computed as the maximum value of the *N_m_* values calculated for all partially resistant mutants. For the lowest risk of *de novo* resistance, the prescribed number of compensatory doses (*N_m_*) should be taken, uninterrupted, immediately after doses are missed. Otherwise the infected cell population may rebound to a high level, which can make further missed doses very risky for resistance.

### Case study: combination therapy of daclatasvir and asunaprevir

To demonstrate the practical applicability of our theory, we consider a recently developed interferon-free combination therapy based on an NS5A inhibitor, daclatasvir, and an NS3 protease inhibitor, asunaprevir (Suzuki, Ikeda et al. 2012). In clinical trials, a large proportion of patients infected with HCV genotype-1b achieved sustained virological response (i.e. viral eradication) when treated with daclatasvir and asunaprevir for 24 weeks, although viral breakthrough and viral relapse occurred in a small fraction of patients (Karino, Toyota et al. 2013; Kosaka, Imamura et al. 2014).

We first consider patients with the wild-type virus at baseline, i.e. the wild-type virus is the dominant strain before treatment. Using the PK/PD data for each drug (Eley, Pasquinelli et al. 2010; Nettles, Gao et al. 2011; Ke, Loverdo et al. 2014) and the resistance profiles data measured for genotype-1b HCV (Fridell, Qiu et al. 2010; Fridell, Wang et al. 2011), we predicted which mutants are potentially fully-resistant to this combination therapy and calculated the values of *N_m_* and Φ*_m_* for each of the partially resistant mutants (Fig. 4A,B) (see Supplementary Materials for more detail). Choosing the highest values of *N_m_* and Φ*_m_* among all the partially resistant mutants allows us to project the overall risk arising from missed doses over the course of treatment, and we found required numbers of compensatory doses were modest and the risk of *de novo* resistance is low (Fig. S2A). To demonstrate that the theoretical approximations represent the full viral dynamics accurately, we simulated a multi-strain viral dynamics model (see Materials and Methods), assuming 1-3 day blocks of consecutive doses are missed randomly within a treatment regimen lasting 24 weeks. The model predicts that relapse of L31M + Y93H or L31W would be observed when overall adherence is less than 90% (Fig. 4C,D). Indeed, the L31M + Y93H mutant has already been detected in one relapse patient in a clinical trial (Karino, Toyota et al. 2013). There is excellent agreement between simulation results and theoretical predictions (based on Eqn.3 and 4) for the number of cells infected by different mutants after 24 weeks of treatment and the cumulative number of cells infected by partially resistant mutants over the treatment period (Fig. 4D and S3).

**Figure 4.**
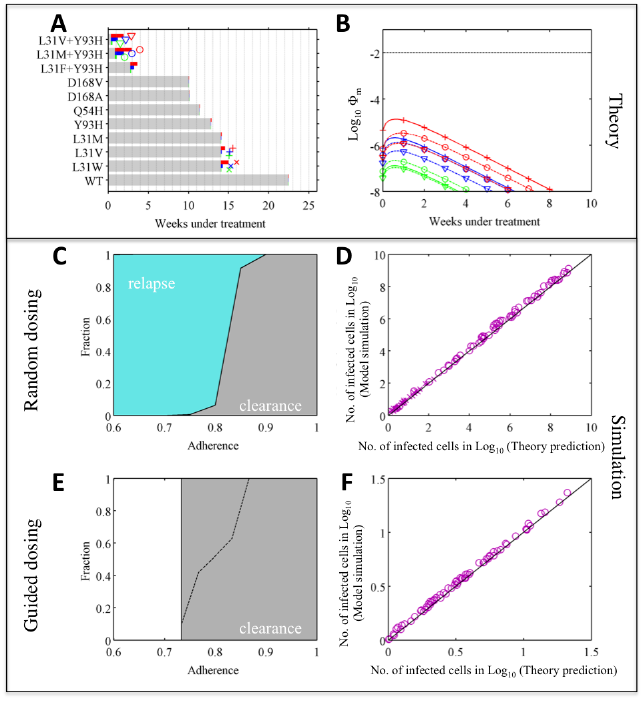
Adaptive treatment strategy improves treatment outcome substantially – a case when the risk of *de novo* resistance is low (with wild-type genotype-1b HCV at baseline). **(A)** Theoretical prediction of the treatment duration needed to eliminate each partially resistant mutant under perfect adherence (gray bar), and the maximum number of additional doses needed to compensate for missing doses, *N_m,max_* (colored bars). Green, blue and red denote results when 1, 2 and 3 consecutive daily doses are missed, respectively. The symbols next to the bars for *N_m,max_*show the type of mutant investigated in panels (B,D,F). **(B)** Theoretical prediction of the risk of *de novo* resistance, Φ*_m_*, over time (as shown in Fig. 3d), for the three mutants with highest risks of generating fully resistant mutants. The dashed black line shows Φ*_m_*=0.01. **(C)** Treatment outcomes 1-3 days of doses are missed randomly. Colored areas denote the fractions of simulations with outcomes of viral relapse without full resistance (light blue) and viral clearance (gray). **(D)** Comparison between theory predictions and simulations of the number of cells infected by different mutants after 24 weeks of treatment. **(E)** Treatment outcomes if adaptive treatment strategy is followed. The area above the black dashed line denotes the fraction of patients where virus is not cleared after 24 weeks’ treatment. After 24 weeks, patients take the prescribed number of make-up doses without missing further doses. White areas denote adherence levels that are not allowed by the adaptive treatment strategy. **(F)** Same comparison as in panel (D) for the guided dosing simulation.

We then simulated outcomes when the doses are guided by the adaptive treatment strategy (*guided dosing*). Because the risk of *de novo* resistance when doses are missed is low, there is no high-risk period for *de novo* resistance in this case (Fig. 4B). If patient dosing is guided, i.e. all the required doses and the extra doses to compensate for the missed doses are taken, the infection can be cleared successfully (Fig. 4E). Again, we find excellent agreement between simulation results and theoretical predictions (Fig. 4F).

Many patients bear the Y93H mutation at baseline and this mutation reduces the genetic barrier to full resistance by one nucleotide(Karino, Toyota et al. 2013). Our theory suggests that reducing the genetic barrier to full resistance will drastically increase the risk of treatment failure. We repeated our analysis for patients with Y93H at baseline, to test how our adaptive treatment strategy works when the risk of resistance is high. As predicted, many more days of treatment are needed to compensate for missed doses, and the risks of generating full resistance *de novo* are high (>0.01) during the first 3 weeks of effective treatment if 2 consecutive doses are missed (or first 4 weeks if 3 doses are missed; Fig. 5A,B and S2B). *De novo* full resistance is likely if doses are missed randomly and adherence is less than 90% (dark red area in Fig. 5C). The predicted number of infected cells agrees well with simulation, except when adherence is very low such that viral load rebounds back close to the pre-treatment level (Fig. 5D and S4–S6). In stark contrast, when doses are guided, the risk of *de novo* resistance becomes much lower (compare Fig. 5C with 5E). Again, for patients who do not clear infection after 24-week treatment, extended periods of treatment as predicted by our theory (using Eqn.3) can clear infection with low risk of resistance. The efficacy of the adaptive treatment strategy is robust across different parameter values (Fig. S7–S12 and Supplementary Materials). Therefore, our treatment strategy can improve clinical outcomes substantially by adjusting on-going treatment based on patient adherence patterns.

**Figure 5.**
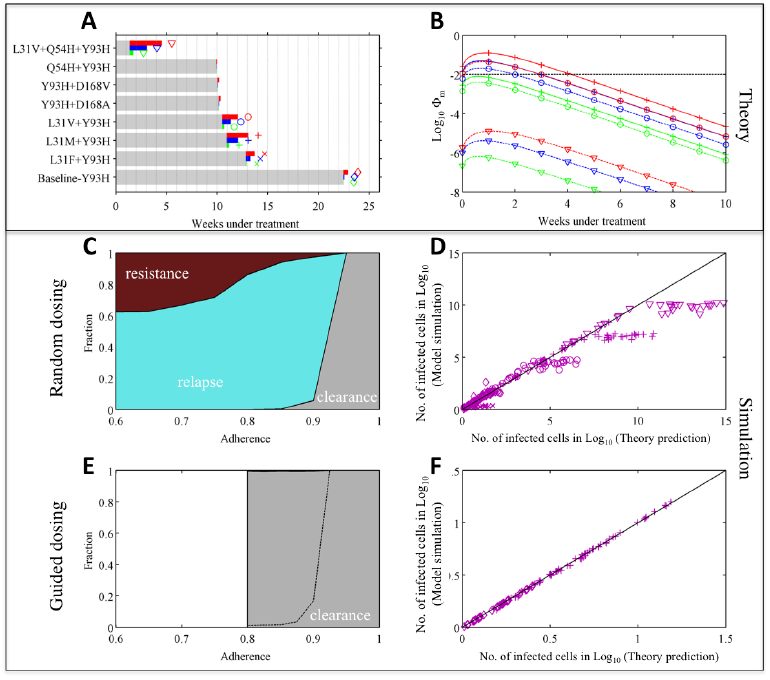
Adaptive treatment strategy prevents *de novo* resistance and improves treatment outcome substantially – a case when the risk of *de novo* resistance is high. Theoretical prediction and simulation for patients with the Y93H mutant virus (genotype-1b) at baseline under combination therapy of daclatasvir and asunaprevir. Thus, the mutants considered here all have the Y93H mutation. The theoretical predictions and simulation results are plotted in the same way as in Fig. 4. Dark red areas in panel (C,E) denote the fraction of patients with *de novo* full resistance to the combination therapy. Note that the fraction of patients with *de novo* resistance in the guided dosing scenario is very small (<0.1%). When doses are guided, so that mutant viral load does not rebound to the pre-treatment level, the theoretical prediction agrees well with simulation as shown in panel (F).

## Discussion

With the prospect of interferon-free combination therapies becoming available to the HCV infected population (Lok, Gardiner et al. 2012; Pol, Ghalib et al. 2012; Scheel and Rice 2013; Thomas 2013; Afdhal, Zeuzem et al. 2014; Feld, Kowdley et al. 2014; Kowdley, Gordon et al. 2014; Zeuzem, Jacobson et al. 2014), there is an urgent need to design treatment strategies that will prevent or delay the development of resistance to DAAs. Extensive laboratory efforts have characterized the PK/PD parameters and mutant resistance profiles of DAAs (Eley, Pasquinelli et al. 2010; Fridell, Wang et al. 2011; Nettles, Gao et al. 2011; McPhee, Friborg et al. 2012; Qi, Olson et al. 2014). In this study, we integrate PK/PD parameters and viral dynamics into a unified framework to assess the impacts of suboptimal treatment adherence on the risk of treatment failure. This framework also enables adaptive management of DAA treatments. Using simulations incorporating PK/PD and resistance profile data collected previously (Fridell, Qiu et al. 2010; Fridell, Wang et al. 2011; Nettles, Gao et al. 2011), we showed that treatment outcomes of combinations therapies of daclatasvir and asunaprevir can be greatly improved by this adaptive treatment strategy, especially when the Y93H mutant is the dominant strain before treatment begins.

For therapies with low genetic barriers to resistance, we have identified a high-risk window period during which *de novo* resistance is likely if doses are missed. Intervention efforts should focus on enhancing patients’ adherence during this period. Additional complementary strategies could further reduce the risk of treatment failure. First, if doses are missed during the high-risk window, the immediate addition of another drug with a different mechanism of action from existing drugs may eliminate any low level of fully resistant mutants that has arisen. Alternatively, a patient could be treated preemptively using additional drugs during the entire high-risk period and switched to fewer drugs afterwards. Our theory also predicts the number of compensatory doses (*N_m_*) needed to compensate for missed doses, in order to eliminate preexisting mutants. Interestingly, clinical trials have shown that adherence levels tend to decrease over time (Weiss, Brau et al. 2009; Lo Re, Teal et al. 2011); we show that more doses are needed to compensate for missed doses that occur later in treatment because of the rebound of target cells. Overall, these results highlight the importance of viral genotype screening and adherence monitoring. While many previous studies have focused on average adherence (Wahl and Nowak 2000; Weiss, Brau et al. 2009; Lo Re, Teal et al. 2011; Evon, Esserman et al. 2013; Gordon, Yoshida et al. 2013; Lieveld, van Vlerken et al. 2013), we emphasize that the timing of the missed doses is also a critical determinant of treatment outcome and the risk of resistance.

There exist substantial heterogeneities among patients owing to variation in HCV genotypes, patient viral loads, death rates of infected cells (Neumann, Lam et al. 1998; Rong, Dahari et al. 2010) and effectiveness of drug penetration (Ke, Loverdo et al. 2014). Our analysis has identified several factors that influence the impact of suboptimal adherence, particularly the rebound rate of target cells under treatment, the half-life of infected cells and the overall viral fitness, *R*_0_. We used the best available estimates of these parameters, but further empirical work is needed. If resistance profiles and viral parameters could be measured directly from a specific patient, then our framework linking these factors could be tailored to give patient-specific guidelines.

Certain model assumptions reflect uncertainties in our current knowledge of HCV infection. First, our prediction about time to viral extinction should be treated cautiously. We predict the time of extinction (as in other models (Snoeck, Chanu et al. 2010; Guedj and Perelson 2011)) by assuming that infected cells decline at a rate set by their death rate, and infection is cleared when the number of infected cells is below one. However, factors such as pressures from the immune system and infections in different tissue compartments may influence the extinction threshold. Furthermore, if DAA treatment causes intracellular viral RNA to decay with negligible replication (Guedj, Dahari et al. 2013), the decline of infected cells may result from a combination of cell recovery and death of infected cells. Indeed, sustained virological response has been observed in clinical trials of DAA combination therapies with shorter durations of treatment (Poordad, Lawitz et al. 2013). Our model can be adjusted easily once the decay dynamics of infected cells are understood better. Second, our model captures the main features of pharmacodynamics and viral dynamics by assuming quasi-equilibrium for viral populations and drug penetration into liver cells. Further work that incorporates detailed intracellular interactions (Guedj, Dahari et al. 2013) and different body compartments may improve model accuracy, once pertinent parameters are measured. However, a more detailed model may become analytically intractable.

This quantitative framework is a step towards developing a tool (for example, see Ref. (Garg, Adhikari et al. 2005)) for clinicians to design combination therapies and adaptively manage treatment regimens to achieve favorable clinical outcomes. It highlights the importance of characterizing resistance profiles of HCV, screening for resistant mutations before treatment, and monitoring adherence patterns during treatment, so that treatment can be designed and adjusted in an evidence-based manner. This framework can be adapted easily to combination therapies based on other DAA candidates, or treatments of other curable diseases without a reservoir.

## Materials and Methods

### HCV model and Viral fitness in the presence of drug, R_eff_(t)

To analyze the dynamics of the virus, we constructed an ordinary differential equation (ODE) model to describe the long-term within-host dynamics of a single HCV strain under drug treatment, based on an established model developed by Neumann *et al*.(Neumann, Lam et al. 1998) (see Supplementary Material). In the model, *ε* represents the proportion by which the therapy reduces viral growth (*ε* is in the range of 0 and 1). Then, the fitness of the virus, *R_eff_*(*t*), is the product of the complement of the therapy’s efficacy (1-*ε*(*t*)), the reproductive number of the virus, *R*_0_, and the availability of target cells, *h*(*t*) (Eqn. 1).

### Average effective viral fitness when m doses are missed, R_ave,m_

To approximate the time-varying viral fitness, *R_eff_*(t), during the period when *m* consecutive doses are missed, we assume that the abundance of target cells stays constant. This is a good approximation, because the length of the period when consecutive doses are missed tends to be short compared to the time scale of target cell rebound. Then the only time-varying quantity in Eqn. 1 is *ε*(*t*). We can calculate the average level of drug inhibition during the period when *m* doses are missed, *ε_ave,m_*, by incorporating parameters for pharmacokinetics and pharmacodynamics (for example, see Wahl and Nowak(Wahl and Nowak 2000)). Then the time-average effective reproductive number, *R_ave,m_*(*t*), for a mutant when *m* consecutive doses are missed starting at time *t* can be expressed as Eqn. 2. In practice, because the precise number of target cells at time *t* is hard to estimate, we can approximate *R_ave,m_* by setting *h*(*t*) = 1, and then *R_ave,m_* becomes *R_ave,m_* (*t*) ≈ (1- *ε_ave,m)_* · *R*_0_. Because *h*(*t*) ≤ 1, this always overestimates the viral fitness and thus is a conservative estimate in terms of guiding treatment.

### The number of compensatory doses needed (N_m_)

To calculate *N_m_* for each mutant, we make the simplifying assumption that the dynamics of the viral populations are at quasi-equilibrium, because changes in the viral populations occur much faster than changes in infected hepatocytes. Then, the dynamics of the number of cells infected by mutant viruses, *I*(*t*), are described by:

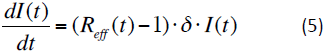

where *δ* is the death rate of infected hepatocytes. If we approximate *R_eff_*(*t*) using the constant *R_ave,m_* for the period when doses are missed, Eqn. 5 can be solved analytically. Then, the number of infected cells after missing *m* consecutive doses starting at time *t*_0_ can be expressed as:

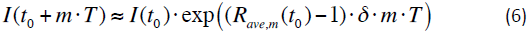

We now consider the situation when *m* consecutive doses are missed, and ask how many uninterrupted doses (compensatory doses) must be taken so that the number of cells infected by the mutant is suppressed to a same number as if the *m* doses had not been missed. We first calculate the number of infected cells if the *m* consecutive doses are taken, i.e. if dosing is optimal:

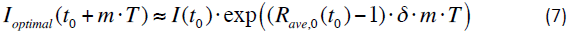

where *I*(*t*_0_) is the number of cells infected by the mutant at time *t*_0_, *R_ave,_*_0_ is the average effective reproductive number of the mutant when all doses are taken, and *T* is the scheduled interval between doses.

We then analyze the situation where a patient skips *m* consecutive doses, starting at time *t*_o_, and then takes *N_m_* compensatory doses immediately afterwards. In this case, assuming the number of target cells does not change much during this period, we can approximate the number of cells infected by the mutant at the end of the *N_m_* doses as:

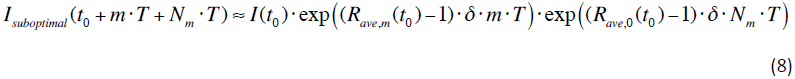

By equating the right hand sides of Eqn. 7 and 8 and solving the equation, we derive the expression for *N_m_*:

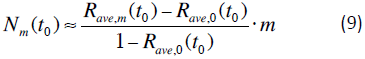

For potent therapies, usually *R_ave_*_,0_ (*t*_0_) ≈ 0. Then we get Eqn. 2.

In the derivation above, we have assumed that the target cell abundance stays constant during the period under consideration. This would be a good approximation if only a few days of doses are missed or if the target cell has already rebounded to the infection-free level. If the abundance of target cells changes considerably during the period under consideration, an alternative, conservative approach would be to assume *h*(*t*) = 1 and take 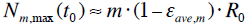 compensatory doses after missing *m* consecutive doses of treatment.

### The number of doses to eradicate a mutant (N_erad_) and the number of cells infected by a mutant (I(t))

One important application of *N_m_* is to predict the number of remaining doses needed to eradicate a mutant, *N_erad_*, in a patient during treatment. This number can be calculated as follows. If adherence is perfect, the number of infected cells declines exponentially at a rate set approximately by the death rate of infected cells, *δ*: *I*(*t*) ≈ *I*_0_ · exp(−δ · *t*), where *I*_0_ is the umber of cells infected by a mutant of interest before treatment. If we assume that a mutant goes extinct if the expected number of infected cells in a patient goes below 1, the number of doses needed to eradicate a mutant before treatment (assuming adherence is perfect), *N_erad,_*_0_, is calculated as: 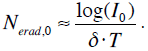.

When doses are missed during treatment, it is clear from the calculation of *N_m_* above that *N_m_*–*m* extra doses of treatment are needed to eradicate the virus. Therefore, if a patient has taken a total of *x* doses and has had *k* instances of missing doses before time *t*, with *m_i_* days of doses missed in the *i*^th^ instance (*i* = 1,2,…,*k*), then the number of remaining doses needed to eradicate the mutant is calculated as:

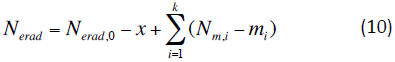

We can use Eqn. 10 to predict the number of cells infected by a mutant as: *I*(*t*) ≈ exp(*δ*·*N*_*erad*_ (*t*)·*T*). In our model, and a patient is cleared of infection when all mutants are driven to extinction. The accuracy of this approximation is shown in Figs. 4D,F and 5D,F.

### The risk of full resistance if doses are missed (Φ_m_)

To calculate the risk of full resistance during the period when *m* doses are missed, we first calculate the number of cells newly infected by a partially resistant mutant when *m* doses are missed, Ω*_m_*(*t*). Again, we use *R_ave,m_*(*t*) to approximate *R_eff_*(*t*), the total number of cells infected by the mutant virus, starting at time *t*. Ω*_m_*(*t*) can be expressed as an integration of new infections during the period of missing doses (according to Eqn.5):

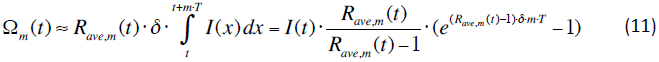

The expected number of target cells that become infected by fully resistant mutant viruses, Φ*_m_*, is a product of the effective mutation rate from the partially resistant mutant to the fully resistant mutant (*μ_eff_*) and the total number of cells infected by the partially resistant mutant (Ω*_m_*): Φ*_m_* (*t*) = *μ_eff_* Ω(*t*), as shown in Eqn.4.

Note that we track the population of newly infected cells to assess the risk of *de novo* generation of full resistance. This assumes implicitly that the fully resistant mutant is selected only when it enters a cell. This is a good assumption for DAAs that act on intracellular stages of the viral life-cycle, such as viral genome replication or assembly. However, in situations where the drug blocks viral entry into the cell, the mutant virus may have a selective advantage for entering a cell. Then the viral population should be tracked instead, but the results presented here still can be applied to drugs that block cell entry by multiplying with a simple scaling factor (Perelson and Nelson 1999).

### Stochastic-deterministic hybrid simulation of multiple strains of HCV

We constructed a simulation model considering the dynamics of the baseline virus and all the potentially partially resistant mutants (see Supplementary Material). This simulation model follows a hybrid approach used previously to simulate the evolutionary dynamics of HIV(Ke and Lloyd-Smith 2012). It considers the dynamics of multiple strains of HCV deterministically (using ODEs) while treating the extinction and mutation processes as stochastic events (see Supplementary Material for detail).

In the simulation, a patient is treated for a total period of 24 weeks. We generate two types of dosing patterns: random dosing and guided dosing. For the random dosing pattern, doses are missed in blocks of 1-3 days at times chosen randomly with equal probability during the treatment period. This probability is set as a constant in each run, but varied across runs such that different overall levels of adherence are generated. In each simulation, we assume that at least one-day treatment is taken immediately after each dose-skipping event (i.e. 1, 2 or 3 consecutive missed doses), to ensure that two dose-skipping events do not occur consecutively (otherwise, longer blocks of doses would be missed than intended). For guided dosing, the procedure is the same as for random dosing, except that we ensure that: 1) doses are always taken during the high-risk window period predicted by our theory, and 2) after the window, a sufficient number of uninterrupted doses (calculated as N*_m_*) are always taken immediately after missing doses, 3) if virus is not eradicated after the 24 weeks treatment period, a patient is treated with an uninterrupted number of doses as predicted by our theory, to ensure eradication of the virus. The outcome of the simulation at the end of the procedure is reported.

## Acknowledgement

We thank Alan Perelson, Paul Fenimore and members in the Lloyd-Smith group and the Sun Group for helpful discussions. This work was supported by National Science Foundation (grant number EF-0928690 to JLS). JLS is grateful for support from the De Logi Chair in Biological Sciences, and from the RAPIDD program of the Science & Technology Directorate, Department of Homeland Security, and the Fogarty International Center, National Institutes of Health.

## Supplementary Text

### 1. HCV model and derivation of viral fitness (R_0_)

We first construct an ordinary differential equation (ODE) model to describe the long-term within-host dynamics of a single HCV strain under drug treatment. This model is based on an established model developed by Neumann *et al*.(Neumann, Lam et al. 1998). It considers the dynamics of the target hepatocytes (H), the infected hepatocytes (I) and the HCV viruses (V). Guedj *et al*. have recently shown that the dynamics of viral loads during the first few days of treatment with direct acting antivirals (DAAs) are better described by a multi-scale model that considers intracellular dynamics of HCV RNAs(Guedj, Dahari et al. 2013). However, since here we are mostly interested in the longer-term dynamics of HCV infection, stretching for weeks or months, the simpler model by Neumann *et al*. is a good approximation.

The ODE model describing the system is:

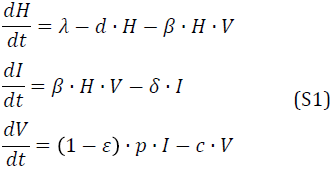

Uninfected target hepatocytes are produced at constant rate, *λ*, and cleared at per capita rate *d*. The infection rate for uninfected hepatocytes is proportional to V, with infection rate constant *β*. The per capita death rate of infected hepatocytes is *δ*. The virions are cleared at per capita rate *c*. In the absence of drug treatment, viruses are produced from infected hepatocytes at rate *p*. Under drug treatment, the production of viruses from infected cells is reduced to rate (1-*ε*)\**p*, where *ε* is the efficacy of the drug of drug treatment.

It has been shown that the number of target hepatocytes increases quickly after initiation of effective treatment (Rong, Dahari et al. 2010). In our model, this rebound rate is determined by the parameters *λ* and *d*, and the number of target hepatocytes in the absence of HCV infection, *H*_0_, is determined by the ratio of this two parameters, i.e. *H*_0_=*λ/d*. We chose values of *λ* and *d* such that the target hepatocytes increase on a similar timescale to the results of Rong *et al*.(Rong, Dahari et al. 2010), while keeping the total number of target hepatocytes constant in the absence of infection. To calculate *R_eff_*(*t*) (in the main text), we set *h*(*t*) as *h*(*t*)=*H*(*t*)/*H*_0_, i.e. the normalized abundance of target cells.

Based on Eqns. S1, we can calculate the reproductive number, *R*_0_, in the absence of drug, as:

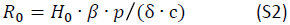

We set *R*_0_=10 in our main analysis, and this choice is in agreement with previous studies(Neumann, Lam et al. 1998; Rong, Dahari et al. 2010).

We assume there are 2*10^11^ hepatocytes in an infected liver(Rong, Dahari et al. 2010). It has been shown that 1%-50% of all hepatocytes are infected in chronically infected patient(Liang, Shilagard et al. 2009; Wieland, Makowska et al. 2014). Thus, we assume that only half of the total hepatocytes can potentially be infected in the absence of treatment, i.e. H_0_=1*10^11^, as in Rong *et al*.(Rong, Dahari et al. 2010). The value of *p* is set such that the reproductive number *R*_0_ of the virus in the absence of drug is 10 (calculated above in Eqn. S2). The parameter values are listed in Table S1.

**Table S1.** Parameter values in the HCV model.

| Parameters | Values | Unit | References |
| --- | --- | --- | --- |
| $\lambda$ | $1.95 \cdot 10^5$ | cells ml<sup>-1</sup> day<sup>-1</sup> | See text |
| $d$ | 0.15 | day<sup>-1</sup> | See text |
| $\beta$ | $8.88 \cdot 10^{-8}$ | ml day<sup>-1</sup> | Rong *et al.* (Rong, Dahari et al. 2010) |
| $\delta$ | 0.15 | day<sup>-1</sup> | Neumann *et al.* (Neumann, Lam et al. 1998) |
| $p$ | 289.76 | day<sup>-1</sup> | See text |
| $c$ | 22.3 | day<sup>-1</sup> | Guedj *et al.* (Guedj, Dahari et al. 2013) |

### 2. Combination therapy of daclatasvir and asunaprevir

#### Pharmacokinetics/Pharmacodynamics

In general, pharmacokinetics of HCV DAAs follow a characteristic pattern: drug concentration increases quickly to a peak level after dosing and then decreases exponentially until the next dose is administered. We use three pharmacokinetic parameters to describe this pattern: the time to reach peak concentration after taking the drug (*τ*), the peak drug concentration (*C_max_*) and the minimum drug concentration before the next treatment (*C_min_*). We assume the active drug concentrations in the liver are related to the drug concentration in the plasma (where data are measured) by a constant ratio *η*. Then, the active tissue concentration of a drug, *C*(*t*), between dosing intervals can be described using the following equation:

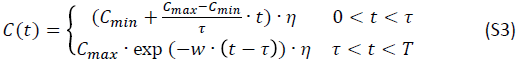

where *C_max_*and *C_min_* are the maximum and minimum drug concentrations in the plasma, *T* is the interval between two consecutive doses, and 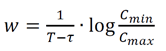. The value of *w* is calculated such that the drug concentration at the beginning is equal to the concentration at the end of a single dose.

The regimen used in clinical trials for the combination therapy of daclatasvir and asunaprevir is 60 mg once daily for daclatasvir and 200 mg twice daily for asunaprevir(Lok, Gardiner et al. 2012; Pol, Ghalib et al. 2012). The value of liver-to-plasma ratio, *η*, for daclatasvir is set as *η* = 0.094 for daclatasvir as shown in recent work(Ke, Loverdo et al. 2014). The value of liver-to-plasma ratio for asunaprevir in human subject is still not clear, although this drug has been shown to have a large liver-to-plasma ratio in animal models(McPhee, Sheaffer et al. 2012). Clinical data from treated patients shows that mutant Q80L + D168 V has EC_90_(the drug concentration suppress 90% production) of 55 nM, and it is resistant to asunaprevir treatment in a patient (PT-29) with trough plasma concentration of 18-33nM(Karino, Toyota et al. 2013). This suggests that the tissue concentration of asunaprevir in the liver is at a similar level to its level in the plasma, and therefore, we have set *η* = 1.0 for asunaprevir. The parameters used for the pharmacokinetics of these two drugs are shown in Table S2.

**Table S2.** Pharmocokinetic parameter values for daclatasvir and asunaprevir treatment used in the simulation model.

| Parameter | Daclatasvir<br>(Nettles, Gao<br>et al. 2011)<br>60mg QD | Asunaprevir<br>(Eley,<br>Pasquinelli et<br>al. 2010)<br>200mg BID |
| --- | --- | --- |
| $C_{max}$ | 1726 ng/ml | 268 ng/ml |
| $C_{min}$ | 255 ng/ml | 35 ng/ml |
| T | 1 day | 0.5 day |
| $\tau$ | 1.5 hour | 3.0 hour |
| $\eta$ | 0.094 | 1 |

Since daclatasvir and asunaprevir act independently on NS5A and NS3 genes, here in the model, we calculate the inhibition of viral growth using Bliss independence(Bliss 1939):

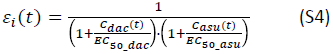

where *C_aac_*(t) and *C_asu_*(t) are the active tissue concentrations of daclatasvir and asunaprevir, respectively, and *EC*_50*_dac*_ and *EC*_50*_asu*_ are the corresponding EC_50_ values of for the viral strain under consideration. The average inhibition during the period when *m* doses are missed, *ε_ave,m_*, is calculated numerically using Eqns. S3 and S4. Another model for pharmacological independence is Loewe independence(Loewe and Muischnek 1926). Changing the model to Loewe independence slightly changes our prediction about the fitness of each mutant under treatment, but does not alter the conclusion of the model.

#### Characterizing preexisting mutants (PMs) and predicting the time needed to eradicate the PMs

We first approximate the equilibrium level of cells infected by the baseline virus. The baseline virus is the viral strain that dominates the population before treatment, i.e. either the wild-type or the Y93H mutant in this study. Its equilibrium abundance before treatment can be derived from the single strain model in Eqn. S1, yielding:

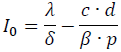

Under effective treatment, the population of infected cells declines exponentially, at a rate set by the half-life of the infected cell (1/*δ*). Then, the time needed to eradicate the baseline virus under perfect adherence can be calculated as:

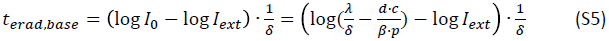

where *I_ext_*is the extinction threshold of infected cells below which the virus goes extinct. In our model, we set I_ext_=1/15000 copy/ml (assuming there are 15L of extracellular fluid(Rong, Dahari et al. 2010)).

If the fitness of a mutant relative to the wild-type is 1-*s_mut_*, then the frequency of resistant mutant virus before treatment can be approximated as(Ribeiro, Bonhoeffer et al. 1998):

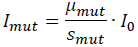

where μ*_mut_* is the mutation rate from the baseline virus to the mutant. Then the time needed to eradicate the mutant virus*, t_erad,mut_*, can be calculated as:

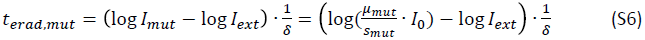

In the model, the mutation rates are set as 2.85*10^-5^ and 1.5*10^-6^ per infection cycle in infected cells for transitions and transversions, respectively (Loverdo et al, unpublished work). A mutant is considered as a preexisting mutant if the number of infected cells calculated for the mutant is above 1 copy in a patient, which corresponds to 1/15,000 copy/ml(Rong, Dahari et al. 2010). If a mutant strain has a higher replicative fitness than the wild-type virus (as measured in the replicon system), we set its relative fitness to 0.99 to ensure that the baseline virus is the dominant strain before treatment. This assumption is required by definition of the baseline strain, and reflects possible differences between fitnesses measured in replicon systems and *in vivo*.

#### Characterizing fully resistant mutants

Since daclatasvir and asunaprevir act independently on different target genes (NS5A and NS3, respectively), we define mutant viruses bearing resistance mutations, i.e. mutants show positive growth under therapy, against both daclatasvir and asunaprevir as potentially fully-resistant to the combination therapy.

To characterize these mutants, we first find mutations that cause resistance, i.e. positive growth, under monotherapies of daclatasvir or asunaprevir, and we assume that daclatasvir and asunaprevir concentrations are the same with the corresponding concentrations in the combination therapy. For each mutant that have been reported to show higher resistance level than the wild-type, we calculated the effective reproductive number under daclatasvir and asunaprevir monotherapies (*R_eff,dac_max_* and *R_eff,asu_max_*, respectively) when the target cell population is at its infection-free level (Table S3 and S4). We find that, if the wild-type virus is the baseline strain before treatment, the preexisting mutants that are resistant to daclatasvir monotherapy are L31M/V+Y93H, and the preexisting mutants resistant to asunaprevir monotherapy are D168A/V (Table S3). Then, combinations of these mutations are potentially fully resistant mutants, e.g. L31V+Y93H+D168 V. If the Y93H mutant virus is the baseline strain before treatment, the preexisting mutants that are resistant to daclatasvir monotherapy are L31M/V+Y93H and L31V+Q54H+Y93H, and the preexisting mutants that are resistant to asunaprevir monotherapy is the same with the case when the wild-type virus is at baseline, i.e. D168A/V (Table S4).

**Table S3.** The resistance profile of genotype 1b mutants when the wild-type virus is at baseline.

| Mutant† | EC<sub>50</sub> to daclatasvir treatment* | EC<sub>50</sub> to asunaprevir treatment* | Relative replication (1-$S_{mut}$) * | $R_{eff,dac\_max}$ | $R_{eff,asu\_max}$ |
| --- | --- | --- | --- | --- | --- |
| **WT** | 0.0026 | 0.86 | 1.0 | 0.0004 | 0.0716 |
| **L31M** | 0.0084 | 0.86 | 0.99** | 0.0011 | 0.0708 |
| **L31V** | 0.0716 | 0.86 | 0.99** | 0.0096 | 0.0708 |
| **L31W** | 0.2100 | 0.86 | 0.99** | 0.0281 | 0.0708 |
| **Q54H** | 0.0032 | 0.86 | 0.83 | 0.0004 | 0.0594 |
| **Y93H** | 0.0621 | 0.86 | 0.27 | 0.0023 | 0.0193 |
| **D168A** | 0.0026 | 109 | 0.37 | 0.0001 | **1.6253** |
| **D168V** | 0.0026 | 241 | 0.29 | 0.0001 | **1.7968** |
| **L31F+Y93H** | 14.87 | 0.86 | 0.29 | 0.4673 | 0.0208 |
| **L31M+Y93H** | 18.23 | 0.86 | 0.70 | **1.3248** | 0.0501 |
| **L31V+Y93H** | 37.94 | 0.86 | 0.50 | **1.5923** | 0.0358 |
| Q54H+Y93H | 0.0243 | 0.86 | 0.22 | 0.0007 | 0.0157 |
| L31V+Q54H+Y93H | 48.74 | 0.86 | 0.99** | 3.6766 | 0.0708 |

† Bold names denote mutants that are preexisting in a patient.
\* Data taken from Fridell et al.(Fridell, Qiu et al. 2010) and McPhee et al.(McPhee, Friborg et al. 2012).
\*\* The fitness values of these mutants relative to the wild-type are set to 0.99, because the values measured in the replicon system were higher than 1 .

**Table S4.** The resistance profile of genotype 1b mutants when the Y93H virus is at baseline.

| Mutant† | EC<sub>50</sub> to daclatasvir treatment* | EC<sub>50</sub> to asunaprevir treatment * | Relative replication (1-$S_{mut}$) * | $R_{eff,dac\_max}$ | $R_{eff,asu\_max}$ |
| --- | --- | --- | --- | --- | --- |
| **Baseline-Y93H\*\*** | 0.0621 | 0.86 | 1.0 | 0.0084 | 0.0716 |
| **L31F+Y93H** | 14.87 | 0.86 | 0.29 | 0.4673 | 0.0208 |
| **L31M+Y93H** | 18.225 | 0.86 | 0.70 | **1.3248** | 0.0501 |
| **L31V+Y93H** | 37.935 | 0.86 | 0.50 | **1.5923** | 0.0358 |
| **Q54H+Y93H** | 0.0243 | 0.86 | 0.22 | 0.0007 | 0.0157 |
| **Y93H+D168A** | 0.0621 | 109 | 0.37 | 0.0031 | **1.6253** |
| **Y93H+D168V** | 0.0621 | 241 | 0.29 | 0.0024 | **1.7968** |
| **L31V+Q54H+Y93H** | 48.735 | 0.86 | 0.99*** | **3.6766** | 0.0708 |

† Bold names denote mutants that are preexisting in a patient.
\* Data taken from Fridell et al.(Fridell, Qiu et al. 2010) and McPhee et al.(McPhee, Friborg et al. 2012).
\*\* We assume that the baseline virus, Y93H mutant, has the highest fitness, set its relative fitness as 1.0.
\*\*\* The fitness value of this mutant relative to the wild-type is set to 0.99, because the value measured in the replicon system is higher than 1.

### 3. Simulation of a hybrid multi-strain model

We construct a simulation model considering the dynamics of the baseline virus and all the preexisting mutants that are shown in Table S3 and S4. This simulation model follows a hybrid approach used previously for simulating HIV evolutionary dynamics(Ke and Lloyd-Smith 2012). The model considers the dynamics of multiple strains of HCV deterministically, using ODEs, while treating the extinction and generation of mutants as stochastic events. The model tracks the population of uninfected and infected hepatocytes. Since the dynamics of viruses are much quicker than those of infected cells, we assume that the virus population is at quasi-equilibrium with respect to the dynamics of the infected cell population: the viral abundance is then given by *V*(*t*) = (1 − *ε*)·*p*·*I*(*t*)/*c*. Then, the ODEs describing the dynamics of the multi-strain system become:

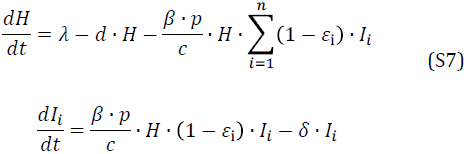

where *H* is the concentration of target hepatocytes, and *I_i_* is the concentration of hepatocytes infected by viral strain *i*.

The mutation process is treated stochastically. During the simulation, the ODEs are first simulated for a fixed time increment (Δ*t* = 0.01 day). At the end of each time increment, we approximate the number of cells newly infected by viruses from cells infected by the i^th^ strain as 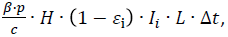 where *L* is the total volume of the liver. Of these newly infected cells, the number of cells in which the infecting viral lineage mutates from the i^th^ strain to the j^th^ strain can be drawn from a binomial distribution with probability of *u*_*ij*_, which is the mutation rate from the i^th^ to the j^th^ strain. We then convert the number to concentration by dividing the number by *L*:

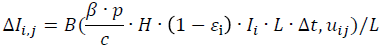

Each time step, the concentration of each strain is updated according to the values of Δ*I_i,j_*. We then check the number of cells infected by each strain. If the number is less than 1 copy per individual, we set it to 0 in the system, i.e. extinction. These procedures are iterated until all infected cells are extinct or simulation time exceeds 24 weeks (for random dosing pattern) or guided dosing period (for guided dosing).

The dosing pattern is generated according to the procedure described in Online Methods. Once the dosing pattern is generated, the drug concentrations are calculated according to Eqn. S3.

### 4. Sensitivity analysis

In the analytical derivations, the parameters that determine the values of N_m_ and Φ_m_ are: the rate at which target hepatocytes become available under treatment (set by the values of parameters *λ* and *d*); the overall viral fitness, *R*_0_; and the clearance rate of infected hepatocytes, *δ*. We performed two rounds of sensitivity analysis, first testing how changes in these parameter values impact the clinical outcomes predicted by our theory, and then testing the robustness of our adaptive treatment strategy to changes in these parameter values.

#### Sensitivity of predicted clinical outcomes to changes in parameter values

##### The overall viral fitness, *R*_0_

The viral fitness parameter, *R*_0_, has several impacts on the predictions of the model. To evaluate the impact of changes in *R*_0_ on the time needed to eradicate the virus, we can substitute the expression of *R*_0_ into Eqn. S5 as:

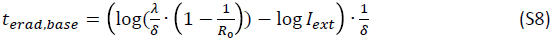

From this equation it can be seen that, if *λ* and *δ* are kept constant, higher R_0_ leads to a higher level of infected hepatocytes before treatment, and thus, a longer time needed to eradicate the virus. However, this increase may be small, because *t_erad,base_* changes in proportion to the logarithm of 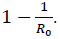.

The value of N_m_ scales linearly with *R*_0_ (Eqn. 2 in the main text). Thus, the higher the value of *R*_0_, the more doses needed to compensate for any missed doses.

The value of Φ_m_ increases almost exponentially with an increase in *R*_0_ (Eqn. 3 in the main text). Therefore the risk of generating fully-resistant mutants increases drastically as *R*_0_ rises (see Fig. S1a,b). This is because the viral population of a mutant increases exponentially if *R_eff,m_* becomes greater than 1, which leads to an exponential increase in the risk of generating fully resistant mutants.

##### The clearance rate of infected hepatocytes, *δ*

The clearance rate of infected hepatocytes, *δ*, influences our theoretical predictions in two ways. First, the time needed to eradicate the virus depends linearly on the inverse of the clearance rate, 1/*δ* (Eqn. S5 and S6). The quicker the clearance rate of infected hepatoctyes, the shorter time needed to eradicate the virus. Second, changes in the value of *δ* affect the risk of generating fully resistant mutants when doses are missed (Eqn. 3 in the main text). If we let the fitness of a virus, *R*_0_, unaffected by changes of *δ*, larger values of *δ* lead to shorter half-lives of infected hepatocytes, and higher risk of generating fully-resistant mutants, because the viral lineage undergoes more rounds of replication during a fixed dosing period (Fig. S1c).

##### The rate at which target hepatocytes become available under treatment

N_m_ is linearly dependent on the level of target hepatocytes (Eqn.2 in the main text), and thus, slower rebound of target hepatocytes would decrease the number of compensatory doses needed if doses are missed before the target hepatocyte population rebounds back to its infection-free level.

The rebound rate of target hepatocytes also impacts Φ_m_, through its influence on the effective viral fitness, *R_eff,m_* (Eqn. 3 in the main text). In our model, the rebound rate is set by the values of parameters *λ* and *d*. If we keep the number of hepatocytes before treatment constant, by keeping the ratio of *λ* over *d* constant, then a slower rebound rate (lower value of both *λ* and *d*) results in a substantially reduced rate of generating *de novo* resistance (Fig. S1d).

#### Robustness of adaptive treatment strategy to variations in parameter values

We first tested the robustness of adaptive treatment strategy to lower or higher values of R_0_ (*R*_0_=5 and *R*_0_=15, as opposed to *R*_0_=10 for our main results). When *R*_0_=5, both the number of compensatory doses (N_m_) and the potential to generate *de novo* resistance (Φ_m_) decrease substantially, as predicted by our model (Fig. S6–S7). This leads to a shorter high-risk window period when mutant Y93H is the baseline strain (Fig. S7) and lower adherence levels are required to eradicate the virus. When *R*_0_=15, we observe the opposite pattern: higher adherence levels are required to eradicate the virus and there is a longer high-risk window period when mutant Y93H is the baseline strain (Fig. S8–S9).

We then tested how well our adaptive treatment strategy works when the half-life of infected hepatocytes is shorter. As shown in Fig. S10 and S11, the time needed to eradicate the virus decreases substantially, to 12 weeks of effective treatment. Our adaptive treatment strategy improves clinical outcome especially when Y93H mutant is at the baseline (Fig. S11).

In general, we find our adaptive treatment strategy is robust against variations in key parameter values. Under all parameter values tested, the adaptive treatment strategy delivers substantially better patient outcomes than random dosing with the same overall adherence levels.

## Supplementary Figures

**Figure S1.**
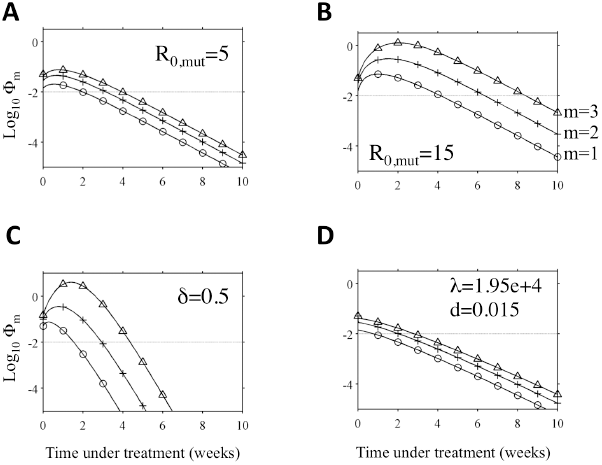
Sensitivity analysis of the risk of *de novo* resistance to variations of key parameter values, R_0,mut_ (panels A,B), *δ* (panel C) and the rate of recovery of target cells upon treatment (panel D). In each panel, the trajectories show how the risk of *de novo* resistance (Log_10_Φ_m_) changes over time if adherence is perfect. Figures are plotted using the same parameter settings as trajectories ‘a’ in Fig. 3 in the main text, except that R_0,mut_=5 in panel A, R_0,mut_=15 in panel B, *δ*=0.5 in panel C and *α*=1.95*10^4^, *d* = 0.015 in panel D. In the main results, i.e. Fig. 3, the parameter values used are R_0,mut_=10, *δ*=0.15, *λ*=1.95*10^5^, *d* = 0.15.

**Figure S2.**
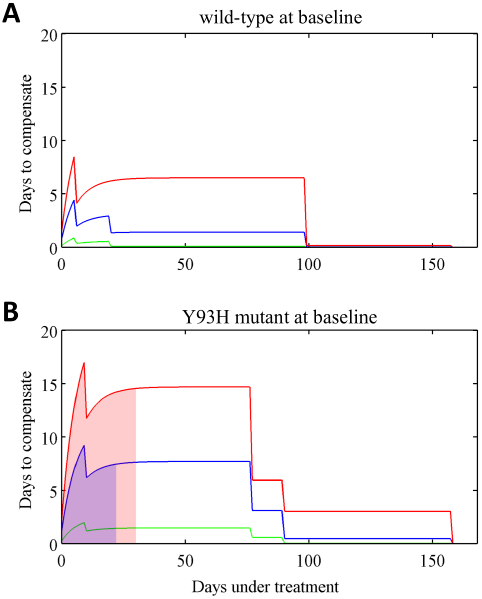
The predicted number of additional days of doses needed to compensate for the first instance of missing 1 (green lines), 2 (blue lines) or 3 (red lines) consecutive days of doses (maximum N_m_ for all partially resistant mutants), and the high-risk window period of *de novo* resistance (shaded area; Φ_m_>0.01 for any of the partially resistant mutants as shown in Fig. 4B). **(A)** Predictions for patients with the wild-type virus at baseline before treatment. The areas below the curves are white, indicating that the risk of *de novo* resistance is always low. **(B)** Predictions for patients with the Y93H mutant virus at baseline. The initial increases of the number of days of compensating doses are due to the increase of the number of target cells upon treatment, and the sudden drops during later periods of treatment are due to the elimination of particular partially resistant mutant lineages. Note that these curves are calculated under the assumption that adherence is perfect except for the 1-3 days of missed doses being considered, i.e. it is a prediction for the first instance of missed doses. For cases where multiple instances of missed doses have occurred, one needs to calculate the values of N_m_ and Φ_m_ for each mutant based on the adherence pattern, and then integrate them together by choosing the highest values of N_m_ and Φ_m_ for those mutants.

**Figure S3.**
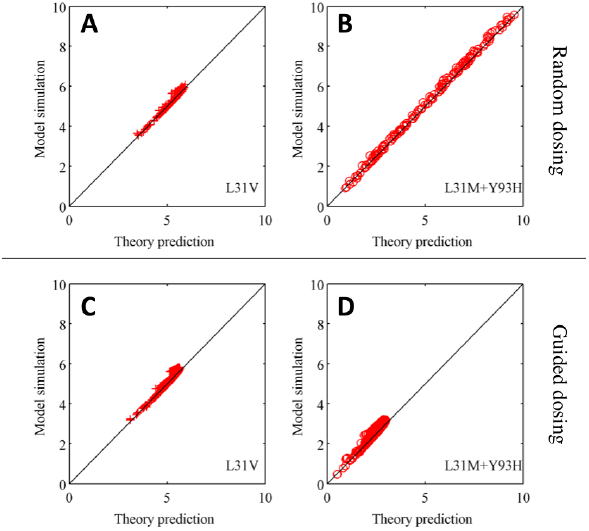
Theory correctly predicts the number of cells infected by two mutants (L31V and L31M + Y93H) generated in the hybrid model simulation when doses are missed randomly (panels A,B) or guided by adaptive treatment theory (panels C,D) for patients with the wild-type virus at baseline. L31V and L31M + Y93H are the two most likely mutants that generate full resistance. The axes are the theory prediction (x-axis) and model simulation (y-axis) of the Log_10_ of the number of mutants, which are calculated as the cumulative numbers of Log_10_ Φ_m_(*t*)/μ*_mut_* for all missed doses.

**Figure S4.**
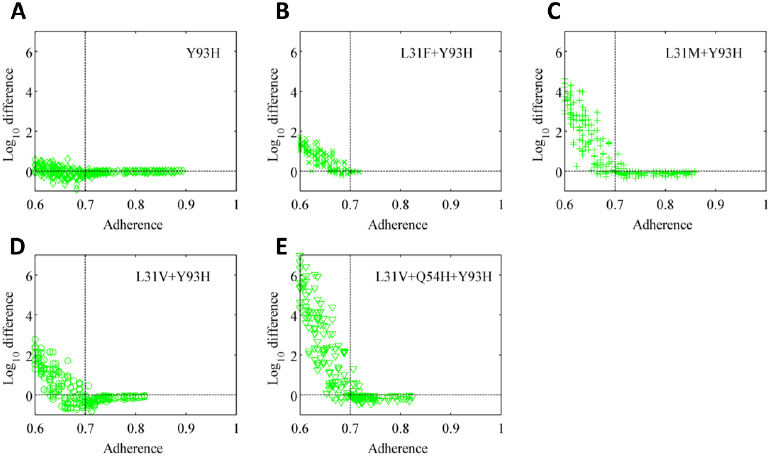
The theory correctly predicts the number of cells infected by mutant viruses at the end of 24-weeks’ treatment in the hybrid model simulation when adherence is greater than 70% (vertical dashed lines) and doses are missed randomly, for patients with the Y93H mutant virus at baseline. The y-axis shows the log_10_ difference between the theory prediction and the model simulation at the end of the 24-weeks’ treatment. Note that when adherence is lower than 70%, the population of infected cells grows to high levels close to the pre-treatment level, where further growth is curtailed by target cell limitation. As a result, the theoretical prediction overestimates the number of cells infected by the mutant virus significantly because we assume the number of target cells is not limited.

**Figure S5.**
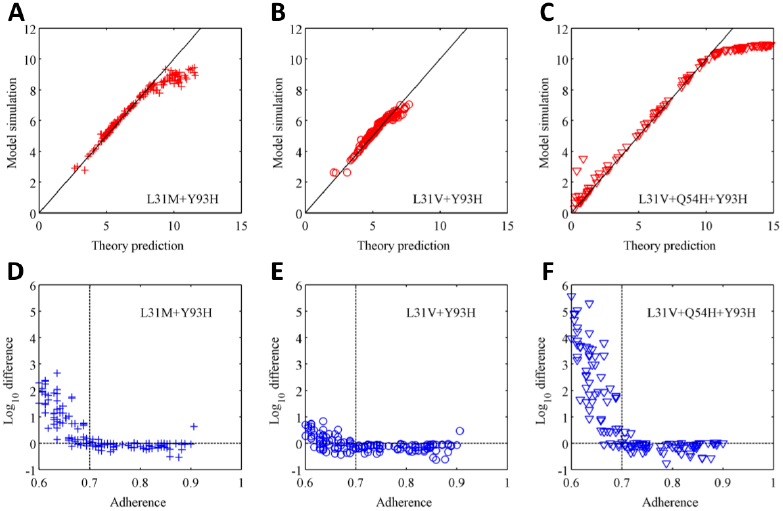
Comparison between theory prediction and simulation of the numbers cells infected by three mutants (L31M + Y93H, L31V + Y93H, L31V + Q54H + Y93H) generated when doses are missed randomly, for patients with the Y93H mutant virus at baseline. **(A,B,C)** The axes are the theory prediction (x-axis) and model simulation (y-axis) of the Log_10_ of the number of cells infected by different mutants, which are calculated as the cumulative numbers of Log_10_ Φ_m_/μ*_mut_* for all missed doses (L31M + Y93H in panel A; L31V + Y93H in panel B; L31V + Q54H + Y93H in panel C). **(D,E,F)** Our theory prediction is accurate for adherence greater than 70%, but overestimates the number of cells infected by the mutant virus significantly when adherence is lower than 70%, for the same reason as explained in the legend of Fig. S4.

**Figure S6.**
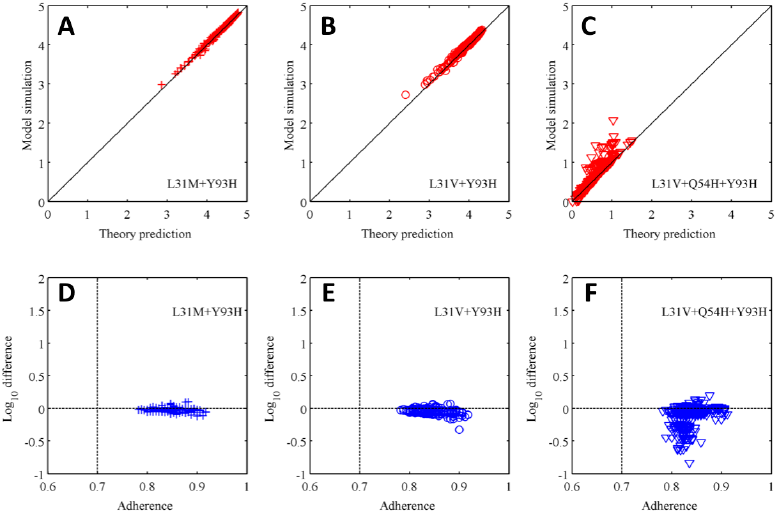
Theory correctly predicts the number of cells infected by three mutants (L31M + Y93H, L31V + Y93H, L31V + Q54H + Y93H) generated in the hybrid model simulation when doses are guided by adaptive treatment theory, for patients with the Y93H virus at baseline. **(A,B,C)** The axes are the theory prediction (x-axis) and model simulation (y-axis) of the Log_10_ of the number of mutants (L31M + Y93H in panel A; L31V + Y93H in panel B; L31V + Q54H + Y93H in panel C), which are calculated as the cumulative numbers of Log_10_ Φ_m_(*t*)/μ*_mut_* for all missed doses. **(D,E,F)** the Log_10_ differences between theory prediction and model simulation as shown in panels (A,B,C). Note that our theory agrees very well for mutants L31M + Y93H and L31V + Y93H. For mutant L31V + Q54H + Y93H, the stochastic extinction and appearance of this mutant generates stochastic deviations of the simulation from theory prediction.

**Figure S7.**
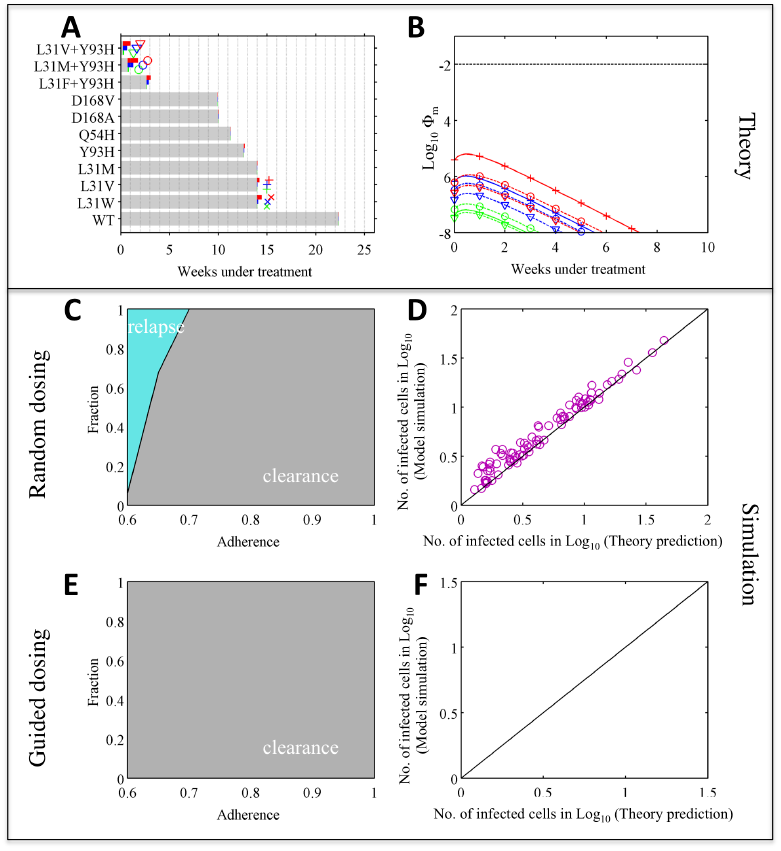
The impact of lower viral fitness (R_0_=5) on treatment outcomes and adaptive treatment strategies of combination therapy with daclatasvir and asunaprevir, with the wild-type virus at baseline. Panels A-F show the same plots as Fig. 4 in the main text, except that the fitness parameter R_0_ for the wild-type virus is assumed to be 5. The treatment outcome improves for all scenarios for this lower viral fitness (compare with Fig. 4 in the main text). Using the adaptive treatment strategy prevents viral relapse and *de novo* resistance if overall adherence is greater than 60% (panel E). Panel F is empty because all patients are cleared of infection after 24 weeks.

**Figure S8.**
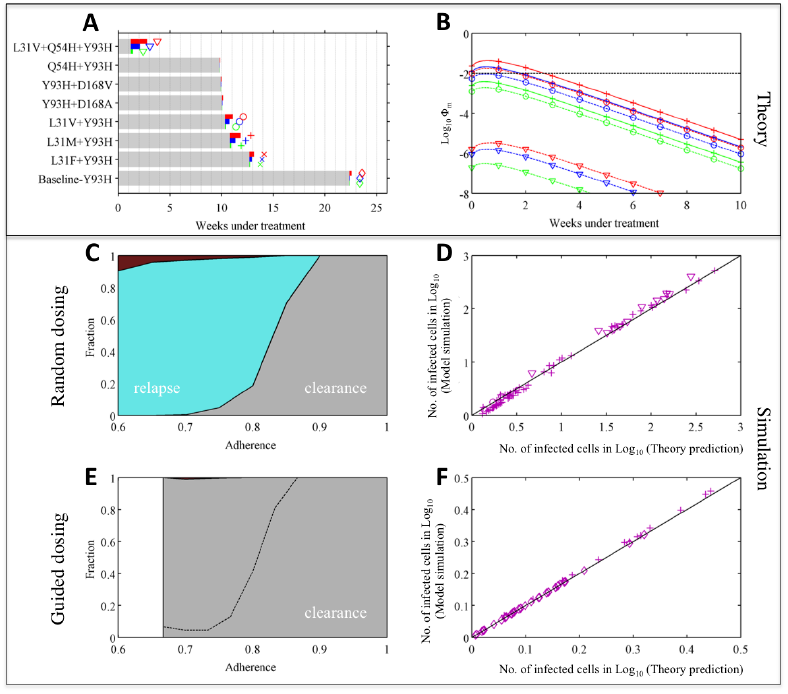
The impact of lower viral fitness (R_0_=5) on treatment outcomes and adaptive treatment strategies of combination therapy with daclatasvir and asunaprevir, with the Y93H virus at baseline. Panels A-F show the same plots as Fig. 5 in the main text, except that the fitness parameter R_0_ for the wild-type virus is assumed to be 5. The treatment outcome improves for all scenarios for this lower viral fitness (compare with Fig. 5 in the main text). Using adaptive treatment strategy reduced the risk of *de novo* resistance (panels E). Our theory correctly predicts the number of infected cells in a patient at the end of 24 weeks’ treatment.

**Figure S9.**
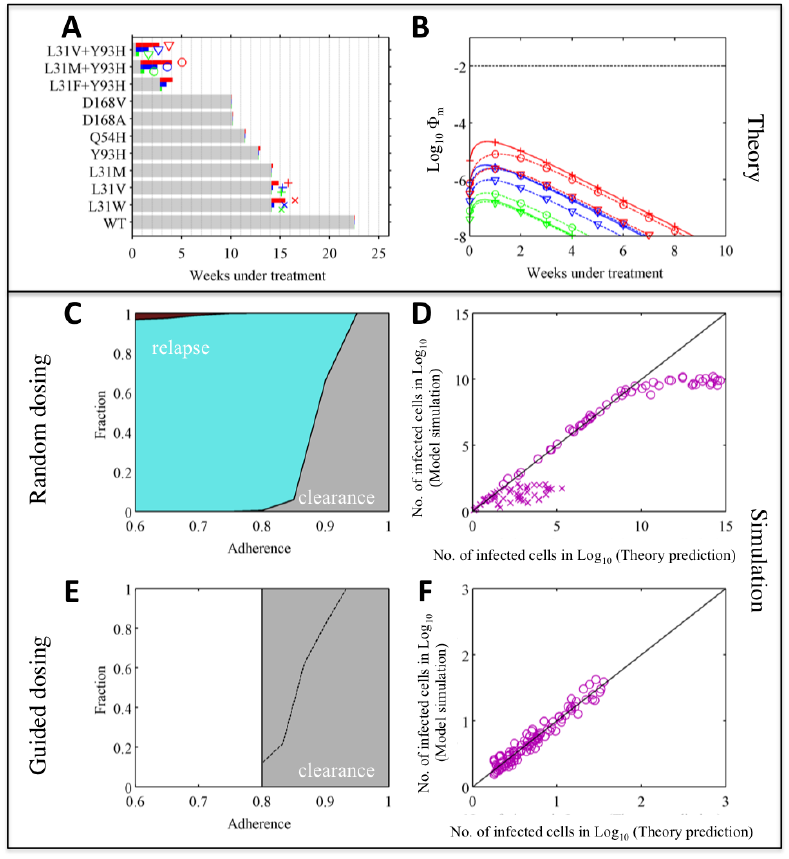
The impact of higher viral fitness (R_0_=15) on treatment outcomes and adaptive treatment strategies of combination therapy with daclatasvir and asunaprevir, with the wild-type virus at baseline. Panels A-F show the same plots as Fig. 4 in the main text, except that the fitness parameter R_0_ for the wild-type virus is assumed to be 15. The treatment outcome improves for all scenarios for this lower viral fitness (compare with Fig. 4 in the main text). Our theory correctly predicts the number of infected cells in a patient at the end of 24 weeks’ treatment.

**Figure S10.**
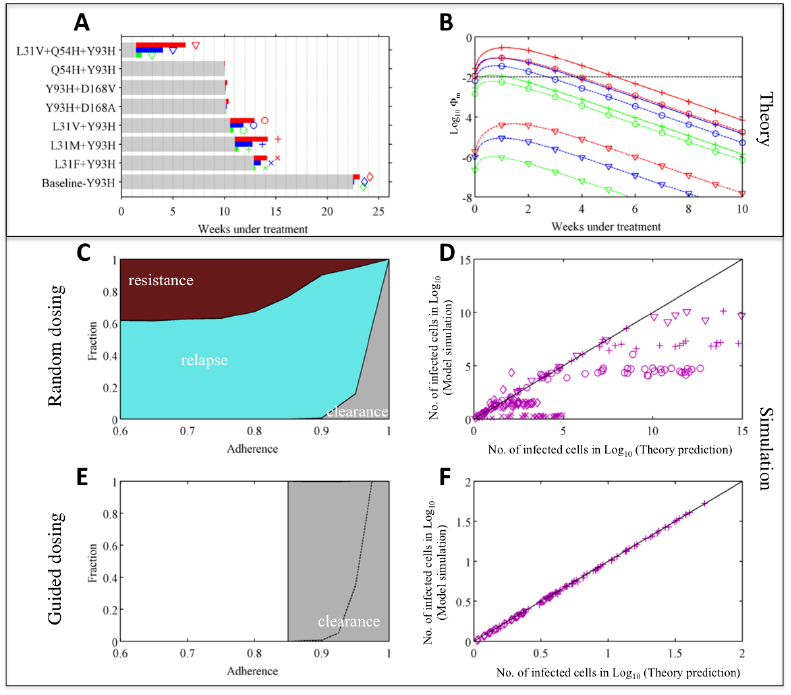
The impact of higher viral fitness (R_0_=15) on the treatment outcomes and adaptive treatment strategies of combination therapy of daclatasvir and asunaprevir with the Y93H virus at baseline. Panels A-F show the same plots with Fig. 5 except that in the analytical derivation and model simulation, the fitness parameter R_0_ for the Y93H mutant virus is assumed to be 15. The risks of viral relapse and *de novo* resistance become higher when the viral fitness, R_0_, is higher. Using adaptive treatment strategy can prevent *de novo* resistance and improve treatment outcomes (panels E and F). Our theory correctly predicts the number of infected cells in a patient at the end of 24 weeks’ treatment when doses are guided (panels F). Our theory does not predict the number of infected cells at the end of treatment well, when doses are missed randomly and the adherence is low. This is because, when adherence is low, the viral load often rebounds back to the pre-treatment level, where it is limited by target cell availability. This phenomenon is not included in our theory, which overestimates the number of viruses as a result.

**Figure S11.**
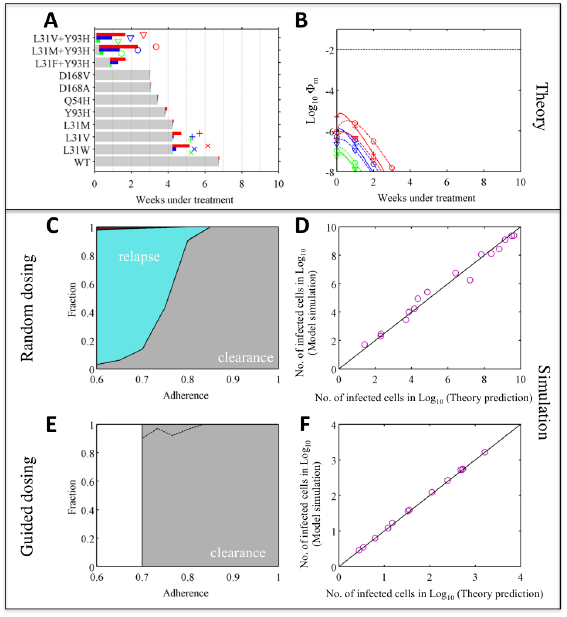
The impact of higher viral clearance rate (*δ* =0.5) on the treatment outcomes and adaptive treatment strategies of combination therapy of daclatasvir and asunaprevir with the wild-type genotype 1b virus at baseline. Panels A-F show the same plots with Fig. 4 except that in the analytical derivation and model simulation, the viral clearance rate, *δ*, is assumed to be 0.5 instead of 0.15 in Fig. 4 (but note that R_0_ for the viruses is kept the same). When the viral clearance rate increases, it takes less time to eradicate the virus from a patient. However, when doses are missed, the population of mutant viruses expands more quickly, because the half-life of the infected cells is shorter and thus it undergoes a higher number of replication generations during the period of missed doses. Using the adaptive treatment strategy can prevent viral relapse and *de novo* resistance and improve treatment outcome (panels E). Our theory correctly predicts the number of infected cells in a patient at the end of 24 weeks’ treatment when doses are guided (panel D).

**Figure S12.**
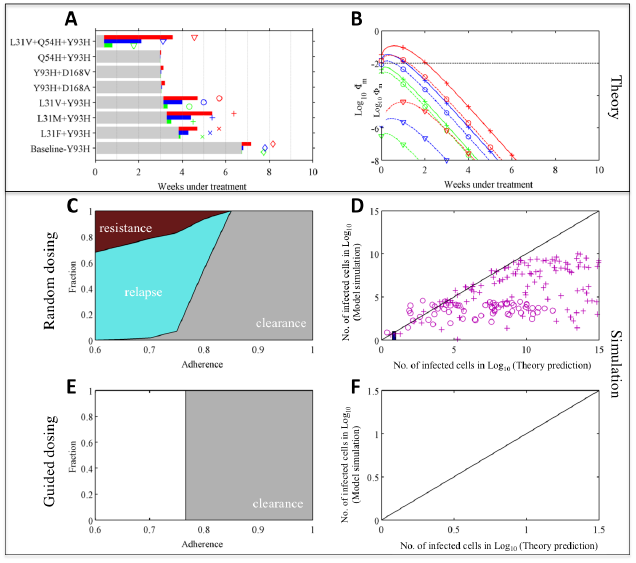
The impact of higher viral clearance rate (*δ* =0.5) on the treatment outcomes and adaptive treatment strategies of combination therapy of daclatasvir and asunaprevir with the Y93H virus at baseline. Panels A-F show the same plots with Fig. 5 except that in the analytical derivation and model simulation, we assume the viral clearance rate, *δ*, is 0.5 instead of 0.15 (but note that R_0_ for the viruses is kept the same). As seen in Fig. S11 for the scenario with the wild-type virus at baseline, it takes less time to eradicate the virus from a patient for this higher viral clearance rate. However, when doses are missed, the population of mutant viruses expands more quickly, increasing the risk of viral relapse and *de novo* resistance. Using adaptive treatment strategy can prevent viral relapse, *de novo* resistance and improve treatment outcome (panels E and F). Our theory does not predict the number of infected cells at the end of treatment well, when doses are missed randomly and adherence is low. This is because, during the time period when doses are missed, the rebound of the viruses is quicker when *δ* is higher (because the viral generation time is shorter). When adherence is low, the viral load often rebounds back to the pre-treatment level, where it is limited by target cell availability. This phenomenon is not included in our theory, which overestimates the number of viruses as a result.

